# Characterization of eyes, photoreceptors and opsins in developmental stages of the chaetognath *Spadella cephaloptera*

**DOI:** 10.1101/871111

**Authors:** Tim Wollesen, Sonia Victoria Rodríguez Monje, Adam Phillip Oel, Detlev Arendt

## Abstract

The phylogenetic position of chaetognaths has been debated for decades, however recently they have been grouped into the Gnathifera, sister taxon to the Lophotrochozoa. Chaetognaths possess photoreceptor cells that are anatomically unique and arranged remarkably different in the eyes of the various species. Studies investigating eye development and underlying gene regulatory networks are so far missing.

In order to gain insights into the development and the molecular toolkit of chaetognath photoreceptors and eyes a new transcriptome of the epibenthic species *Spadella cephaloptera* was searched for opsins. Our screen revealed single-copies of *xenopsin* and *peropsin* and gene expression analyses demonstrated that only *xenopsin* is expressed in photoreceptor cells of the developing lateral eyes. Adults likewise exhibit two *xenopsin*+ photoreceptor cells in each of their lateral eyes. Beyond that, a single *cryptochrome* gene was uncovered and found co-expressed with *xenopsin* in some photoreceptor cells of the lateral developing eye. In addition, it is co-expressed with *peropsin* in the cerebral ganglia, a condition reminiscent of a non-visual photoreceptive zone in the apical nervous system of the annelid *Platynereis dumerilii* that performs circadian entrainment and melatonin release. *Cryptochrome* expression was also detected in cells of the corona ciliata, a circular organ in the posterior dorsal head region that has been attributed several functions arguing for an involvement of this organ in circadian entrainment. Our study demonstrates the importance to investigate representatives of the Gnathifera, a clade that has been neglected with respect to developmental studies and that might contribute to unravel the evolution of spiralian and bilaterian body plans.

## Introduction

Darwin was puzzled by the organizational principles of complex eyes and admitted that it was difficult to view these organs as products of natural selection (Darwin 1859). Ever since then scientists have been intrigued by the organization and evolution of eyes which were assumed to be lost and gained multiple times independently from each other (Salvini-Plawen and Mayr 1977). There is, however, evidence that simple cup-shaped eyes with photoreceptor cells and shading pigments already existed in the last common bilaterian ancestor (Arendt et al. 2004). While rhabdomeric photoreceptors store photopigments (opsins) in expanded surface areas called microvilli, extended cilia are present in ciliary photoreceptors for the same purpose (Arendt et al. 2004). For a long time, the task of vision was thought to be carried out exclusively by rhabdomeric photoreceptors in invertebrates, and exclusively by ciliary photoreceptors in vertebrates (Eakin 1982). Recent studies, however, showed that photoreceptors may also co-express different types of opsins and that distinguishing photoreceptors on morphological grounds is not as clear-cut as previously appreciated (Arendt 2017; Vöcking et al. 2017). Comparative developmental studies on molecular and morphological traits of photoreceptors have shaped an overview on the organization of eyes and photoreceptors and the presence of opsins in diverse bilaterian evolutionary lineages (Arendt and Wittbrodt 2001; Randel et al. 2013; Wollesen et al. 2019). Together, available work now suggests that the last common bilaterian ancestor already possessed several opsins such as a C-opsin, a canonical R-opsin, a non-canonical R-opsin, a Go-opsin, a Neuropsin, and a Retinal pigment epithelium-retinal G protein-coupled receptor/ Peropsin/ Retinochrome (Ramirez et al. 2016).

In contrast to deuterostomes and ecdysozoans, little is known about gene regulatory networks underlying photoreceptor and eye formation in spiralians, except for a few expression studies on mollusks (Vöcking et al. 2015, 2017; Wollesen et al. 2019), annelids (Arendt et al. 2004), platyhelminths (Rawlinson et al. 2019), and brachiopods (Passamaneck and Martindale 2013). The Gnathifera, i.e. sister taxon to the Lophotrochozoa, comprise Chaetognatha, Rotifera, Gnathostomulida, and Micrognathozoa and have not yet been investigated with respect to genes underlying photoreceptor formation and function (Figure 1; Marlétaz et al. 2019). Chaetognaths, commonly known as arrow worms, are a rather small taxon of marine, torpedo-shaped coelomic animals with horizontally projecting fins, and cuticular grasping spines to catch prey (Shinn 1997). As a major planktonic component without respiratory or circulatory systems, chaetognaths exhibit traits reminiscent of deuterostomes (e.g. aspects of gastrulation process) as well as protostomes (nervous system development). Hence their phylogenetic position has been contentious for a long time until phylogenetic analyses recently placed them within the Gnathifera (summarized by Harzsch et al. 2015; Marlétaz et al. 2019).

**Figure 1.**
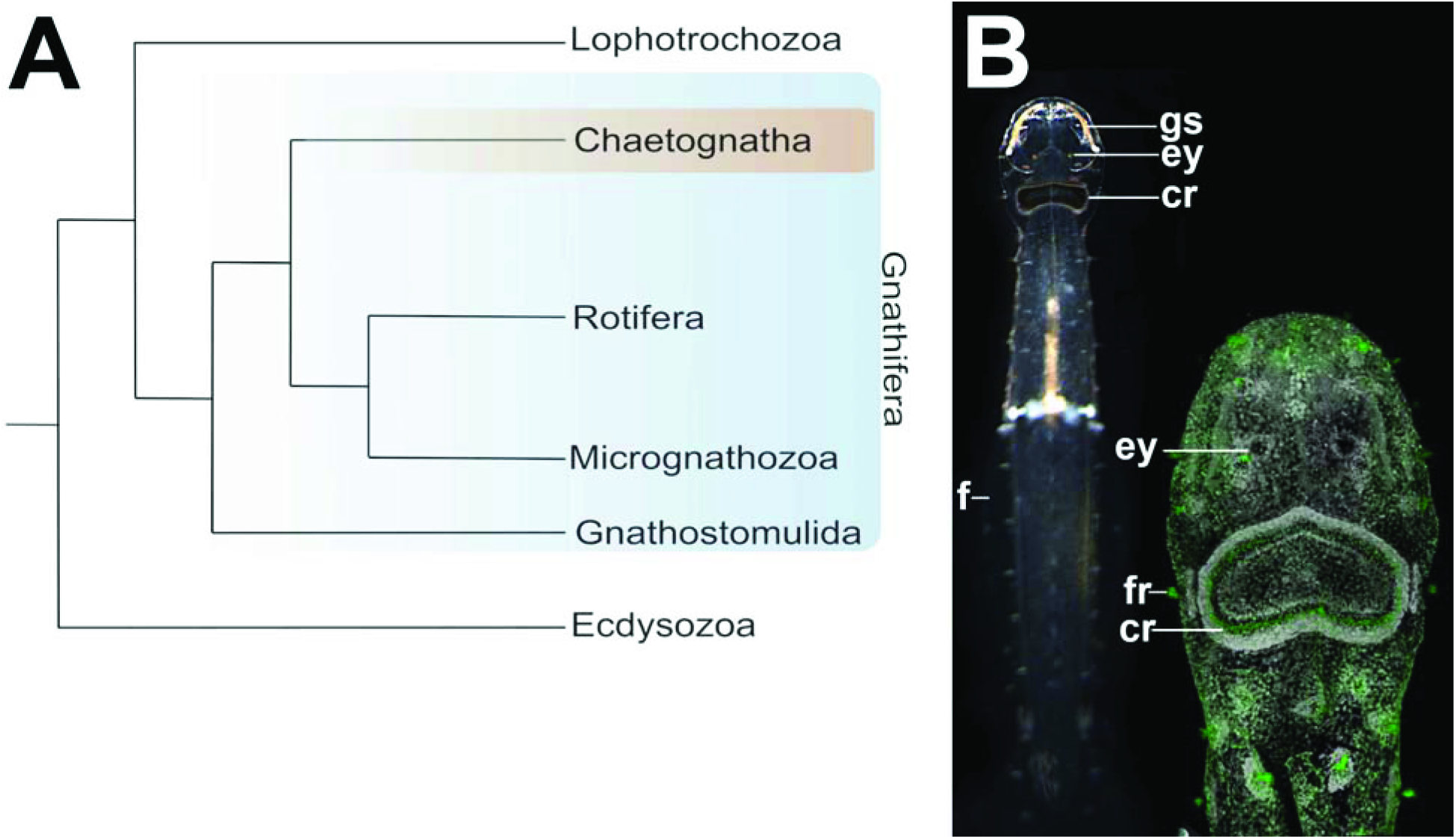
Chaetognaths and their phylogenetic position. **A:** Chaetognatha belongs to the Gnathifera, a taxon being sister to the Lophotrochozoa. The latter includes taxa such as Mollusca, Annelida or platyhelminths and composes together with Gnathifera the Spiralia *sensu* Marlétaz et al. 2019. **B:** An adult of the chaetognath *Spadella cephaloptera* (dorsal view and anterior faces up; individual ∼7 mm in length) (left). Confocal laserscan of the head and anterior trunk region of an adult *S. cephaloptera* (green: tyrosinated tubulin; grey: cell nuclei/ DAPI). As torpedo-shaped predators chaetognaths are equipped with cuticular grasping spines (gs), and horizontally extending fins (f). As sensory organs they possess a pair of compound eyes (ey), numerous fence receptors, and a ring of ciliated cells (‘corona ciliata’ (cr)).

Most chaetognath species possesses a pair of subepidermal eyes that is encapsulated by the extracellular matrix and sheath cells on the dorsal side of the head and that innervates the cerebral ganglia via optic nerves (Fig. 1B; Eakin and Westfall 1964; Goto and Yoshida 1984). Chaetognath photoreceptors within these eyes may be orientated either in an inverted or an everted fashion (Goto and Yoshida 1984). The inverted eye is a spherical dorso-ventrally flattened structure with a pigment cell in the center. Photoreceptor cells are composed of a distal ciliated photoreceptive segment that connects via a conical body with refractive property to the proximal portion (unique to chaetognaths) of the cell ending in an axon (Eakin and Westfall 1964). A single eye may be composed of 70-600 photoreceptor cells (Goto and Yoshida 1984). Photoreceptor cells are arranged around a pigment cell with their distal ciliary photoreceptive segment facing to the center of the eye (‘inverted’) and the axon on the proximal side (Eakin and Westfall 1964; Goto and Yoshida 1984). In ‘everted’ eyes the distal photoreceptive process of each photoreceptor cell points into the periphery and in most of the species investigated more than one pigment cell is present (Goto and Yoshida 1984). Besides their eyes chaetognaths are also equipped with several other sensory organs such as the ciliary fence receptors or the corona ciliata, a ring of ciliated and unciliated cells that is located in the dorsoposterior head region (Müller et al. 2014). Various roles that have been attributed to the corona ciliata include excretory, secretory, or (chemo)sensory function. Ciliary fence receptors have been shown to detect hydrodynamic stimuli and react to close-range mechanosensory input (attack or escape movements) (Horridge and Boulton 1967, Feigenbaum and Reeve 1977, Feigenbaum and Maris 1984, Bone and Goto 1991).

Compared to other bilaterians the inverted photoreceptors of chaetognaths exhibit a unique anatomy with a distal ciliated process connected via a conical body to a cell body with a brush of microvilli, which sends out an axon (Eaking and Westfall 1984). Until now it is, however, unclear which type of opsin is expressed in the chaetognath photoreceptors and in which compartments opsins are expressed, i.e. only in the distal ciliated compartment or also in the microvillar brush? Evidence from histochemistry and peak spectral analysis suggests that a rhodopsin-like pigment is present in the distal processes of the photoreceptors of *Paraspadella gotoi* (Sweatt and Forward 1985; Goto and Yoshida 1988).

In the present study we investigated the presence of opsins and other light-sensitive proteins in developmental stages and adults of the epibenthic chaetognath *Spadella cephaloptera*. This species possesses inverted photoreceptors and is probably the best investigated chaetognath with respect to its nervous and sensory systems (summarized in Harzsch et al. 2015). To this end, we searched for homologs of opsins and other light-sensitive proteins and identified single-copy orthologs of *xenopsin* and *peropsin* as well as *cryoptochrome. Peropsin* orthologs have been found in all major bilaterian groups, while *xenopsin* is only present in Spiralia and possibly in Cnidaria, and may thus be part of the bilaterian group pattern (Sun et al. 1997; Ramirez et al. 2016). *Cryptochrome* is a key player of the circadian system in animals and plays a role in light sensing as well as entrainment of the circadian oscillator (Emery et al. 1998; Thresher et al. 1998; Mei and Dvornyk 2015). In our study we demonstrate that *xenopsin* is probably the only opsin expressed in the chaetognath eyes, in combination with no other opsin. However, we also show that another light-sensitive protein, *cryptochrome*, is expressed in the eyes as well.

## Results

### Phylogenetic and sequence analysis

We identified single-copy orthologs of *xenopsin* and *peropsin* in the transcriptome of *Spadella cephaloptera*. Their predicted protein sequences cluster well with their bilaterian orthologs in our phylogenetic analysis (Fig. S1). Compared to other bilaterians, the characteristic ‘NPXXY’ motif and tripeptide (‘NXQ’) for G-protein activation of the c-terminal Sce-Xenopsin and the Sce-Peropsin contain slightly different residues, i.e. NPLVV + SAR in Sce-Peropsin and DPILY + NKR in Sce-Xenopsin (Passamaneck et al. 2011; Vöcking et al. 2017). The aberrant Sce-Xenopsin residues are identical to those of the Xenopsin found in the chaetognath *Pterosagitta draco* (Rawlinson et al. 2019). Sce-Xenopsin and Sce-Peropsin exhibit both the highly conserved lysine in the retinal binding domain that constitutes the Schiff base with the retinal chromophore forming the photopigment. Sce-Cryptochrome clusters with other type-1 cryptochromes (Figure S2).

### Gene expression and anatomy of the developing chaetognath eyes

Cells contributing to the eye anlagen of early encapsulated embryos of *Spadella cephaloptera* are not lined up next to each other and no pigment cell was observed, however, some photoreceptor cells already express *xenopsin* (Figs. 2A; 3A-C). No *peropsin+* and *cryoptochrome*+ cells are present in these early embryos (not shown). The eyes of young hatchlings are composed of cells that are lined up next to each other and form a spherical shape that is partially filled with cells (Fig. 2B). In hatchlings several *xenopsin+* photoreceptor cells are located in the proximal and distal lateral regions but not in the anterior and posterior regions of the eyes (Fig. 4A-D). During this stage *peropsin* is expressed in the cerebral ganglia and the perikaryal layers of the ventral nerve cord but not in the eyes (Fig. 5A-D). *Cryptochrome* is expressed in photoreceptor cells that are located in the proximal and distal lateral portions of the eyes, in addition to the cerebral ganglia, the corona ciliata, and cells of the outer perikaryal layer of the ventral nerve cord (Fig. 6A-D). The adult eyes are spherical in shape and composed of photoreceptors with distal photoreceptive segments that are arranged into four tightly packed domains (Fig. 2C-D). These four domains share and are separated by a single pigment cell: the two lateral domains are located dorsally, while the anterior (3) and posterior domains (4) are situated more ventrally (Fig. 2C-E). The optic nerves connect the eyes with cerebral ganglia (Fig. 2C). *Xenopsin* is only expressed in each of two photoreceptor cells of their distolateral eye and the region ventrally to the eye (Fig. 7A-D). Adults express *peropsin* in the region antero-ventrally to each eye and faintly in the corona ciliata (Fig. 8A-D). In two-week-old juveniles, *cryptochrome* is expressed in cells of the cerebral ganglia, in cells of the corona ciliata, the ventral nerve cord, and in few cells of the dorsal trunk epidermis (Fig. 9A). In addition, *cryptochrome* is expressed anteroventral to the eyes in each two adjacent cell somata, and in two cell somata posteroventrally to the retrocerebral pore (Fig. 9C). Adults express *cryptochrome* in the same regions but not in the dorsal epidermis (Fig. 9C-D). An additional *cryptochrome*+ cell is located in the region between both above-mentioned expression domains antero-ventrally to the eyes and in the ventral nerve cord (Fig. 9B-D).

**Figure 2.**
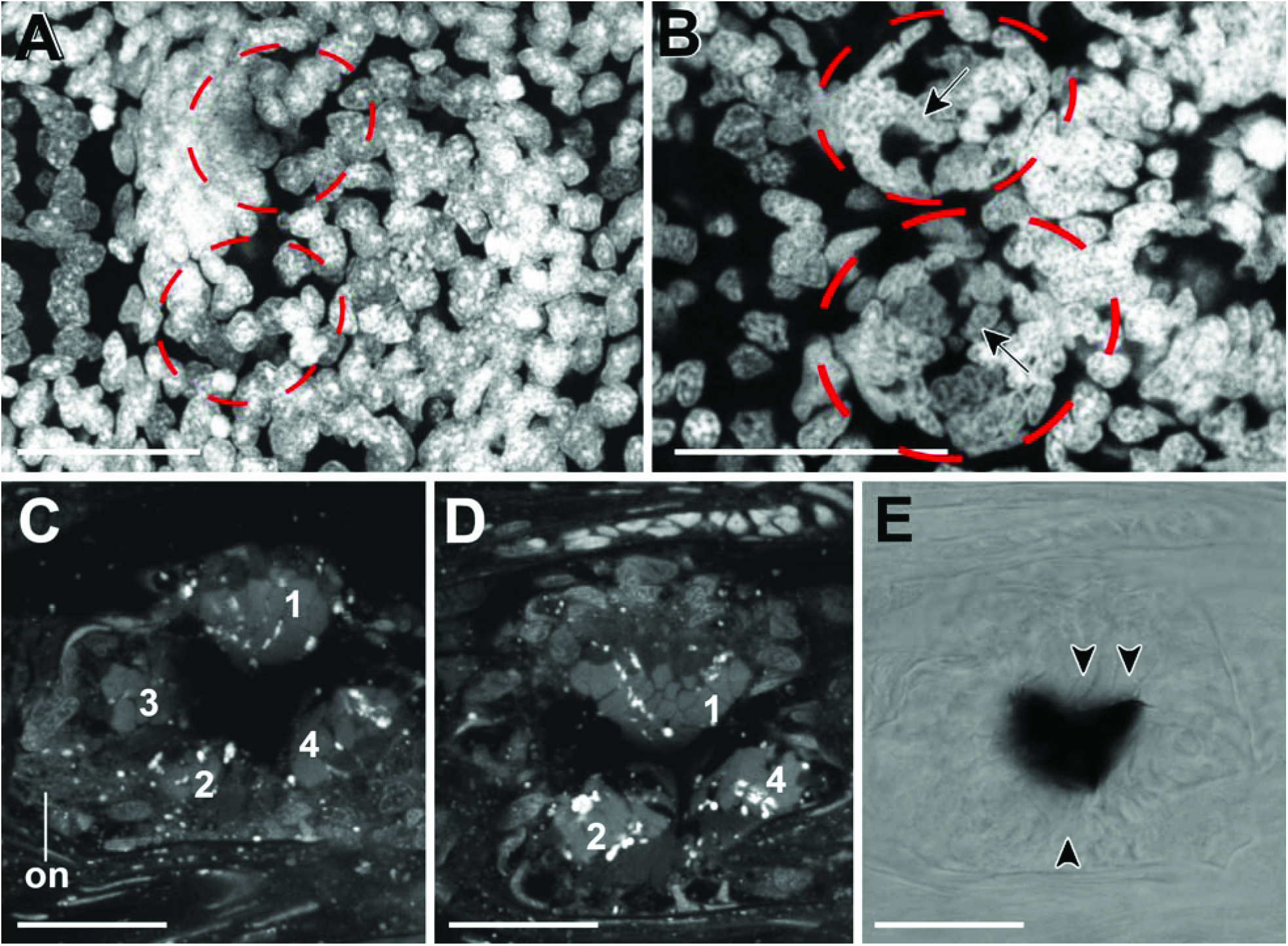
The developing eye of *Spadella cephaloptera*. Dorsal views, anterior to the left. Cell nuclei (DAPI) staining except E (bright field image). **A:** The eyes (encircled) of embryos still surrounded by an egg capsule appear disorganized with cells not arranged in a circular fashion. **B:** In contrast to earlier embryos recently hatched individuals exhibit eyes (encircled) with spherical shape, however, individual cells are still not arranged as seen in subsequent stages (arrow). Eyes do not exhibit pigment cells. **C & D:** In adult *Spadella cephaloptera* the distal segments of photoreceptors are tightly packed into four packages (1-4). While packages 1 & 2 are located more dorsally, packages 3 & 4 are located more ventrally. The optic nerve (on) connects the eye with the cerebral ganglia (not shown). **E:** The pigment cell (black) is surrounded by photoreceptors (arrowheads). Scale bars: 20 µm.

**Figure 3.**
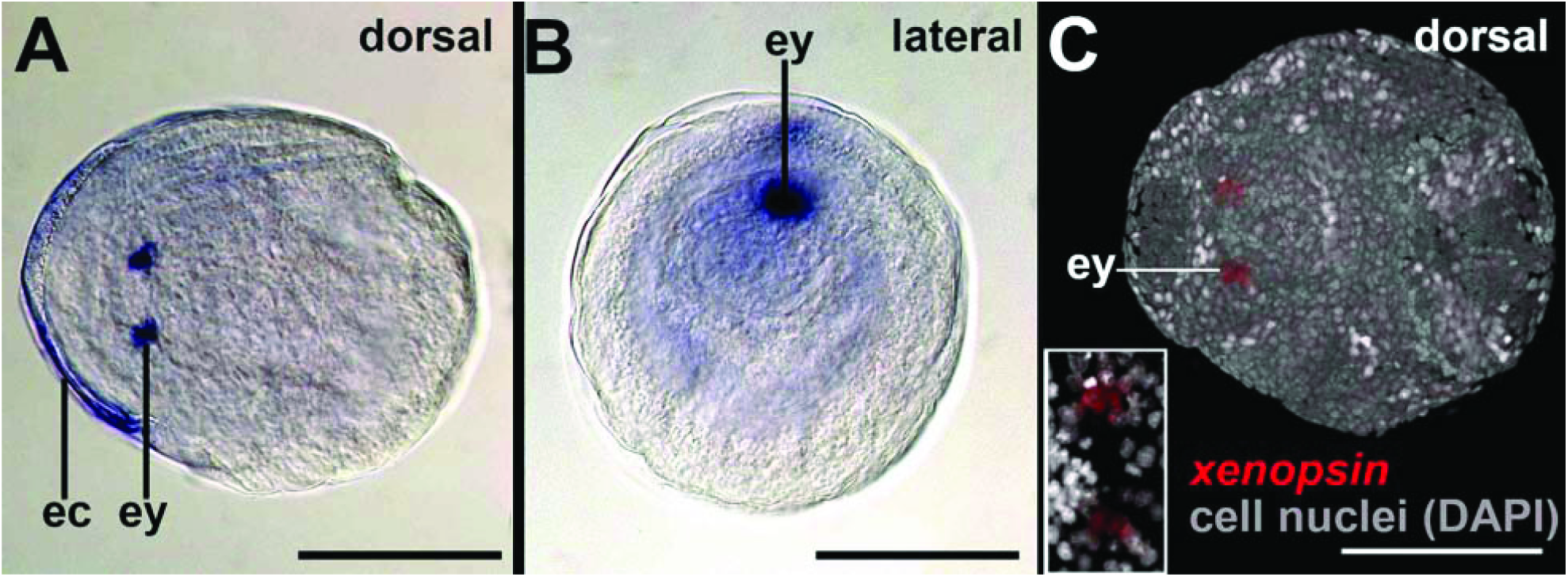
*Xenopsin* expression in early encapsulated embryos of *Spadella cephaloptera*. Whole-mount *in situ* hybridization, anterior to the left. **A:** *Sce-xenopsin+* photoreceptor cells in the eyes (ey) of embryos inside their egg capsule (ec). The egg capsule is unspecifically stained. **B:** The curled-up embryo expresses *sce-xenopsin* in the photoreceptors of its eyes. **C**. Photoreceptor cells are not arranged in a circular pattern highlighted by this confocal laserscanning scan. Scale bars: 150 µm.

**Figure 4.**
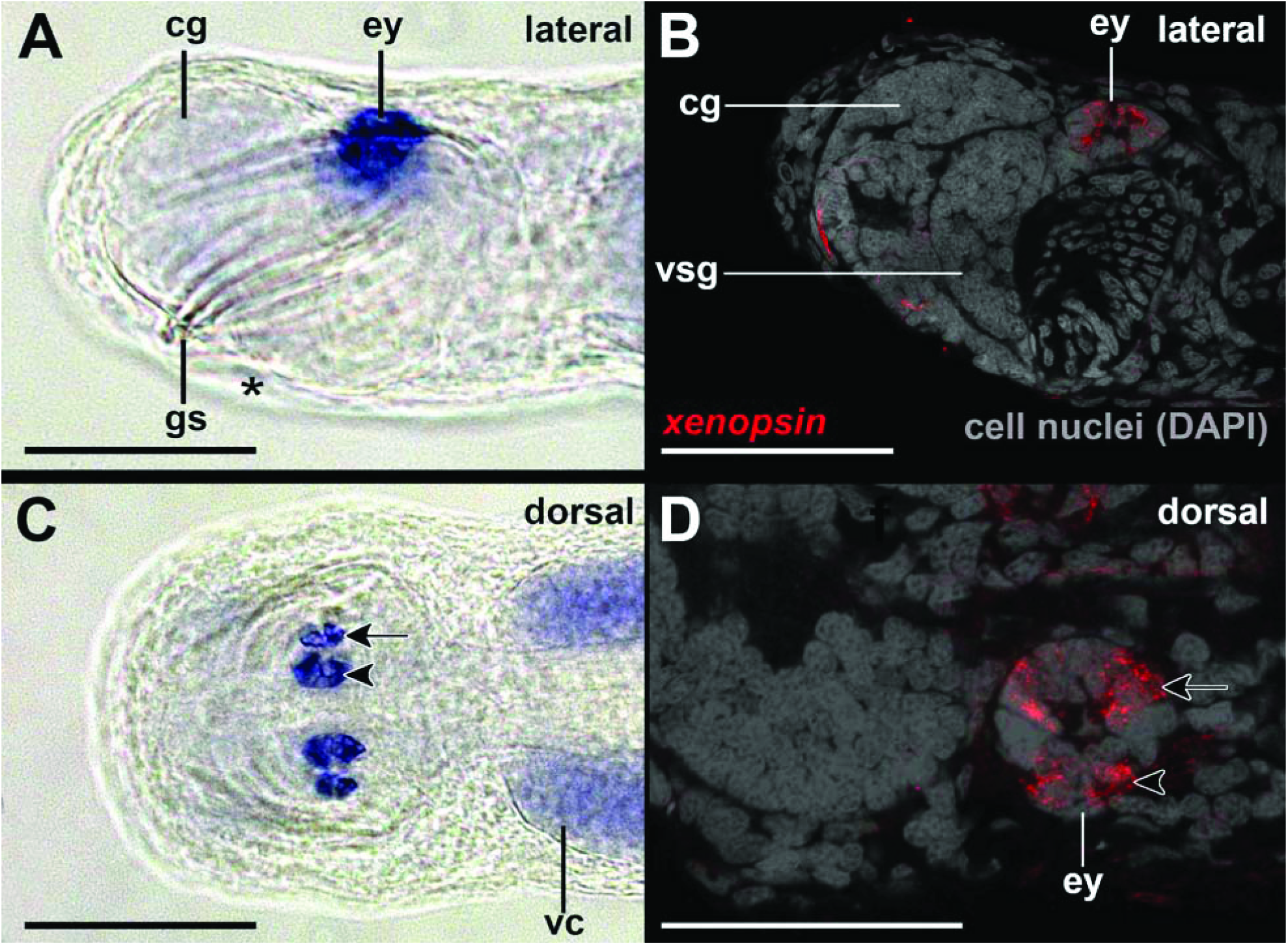
*Xenopsin* expression in early hatchling juveniles of *Spadella cephaloptera.* Whole-mount *in situ* hybridization of the head region, anterior to left. **A:** *Sce-xenopsin+* photoreceptor cells are located in the lateral regions of the eyes (ey) as also highlighted by confocal laserscanning microscopy in **B. C:** *Xenopsin*-expressing photoreceptor cells in the eyes with a close-up by confocal laserscanning microscopy of the left eye in **D**. Abbreviations: cg, cerebral ganglion; gs, grasping spines; vc, ventral nerve cord; vsg, vestibular ganglion. Scale bars: 100 µm (except D: 50 µm).

**Figure 5.**
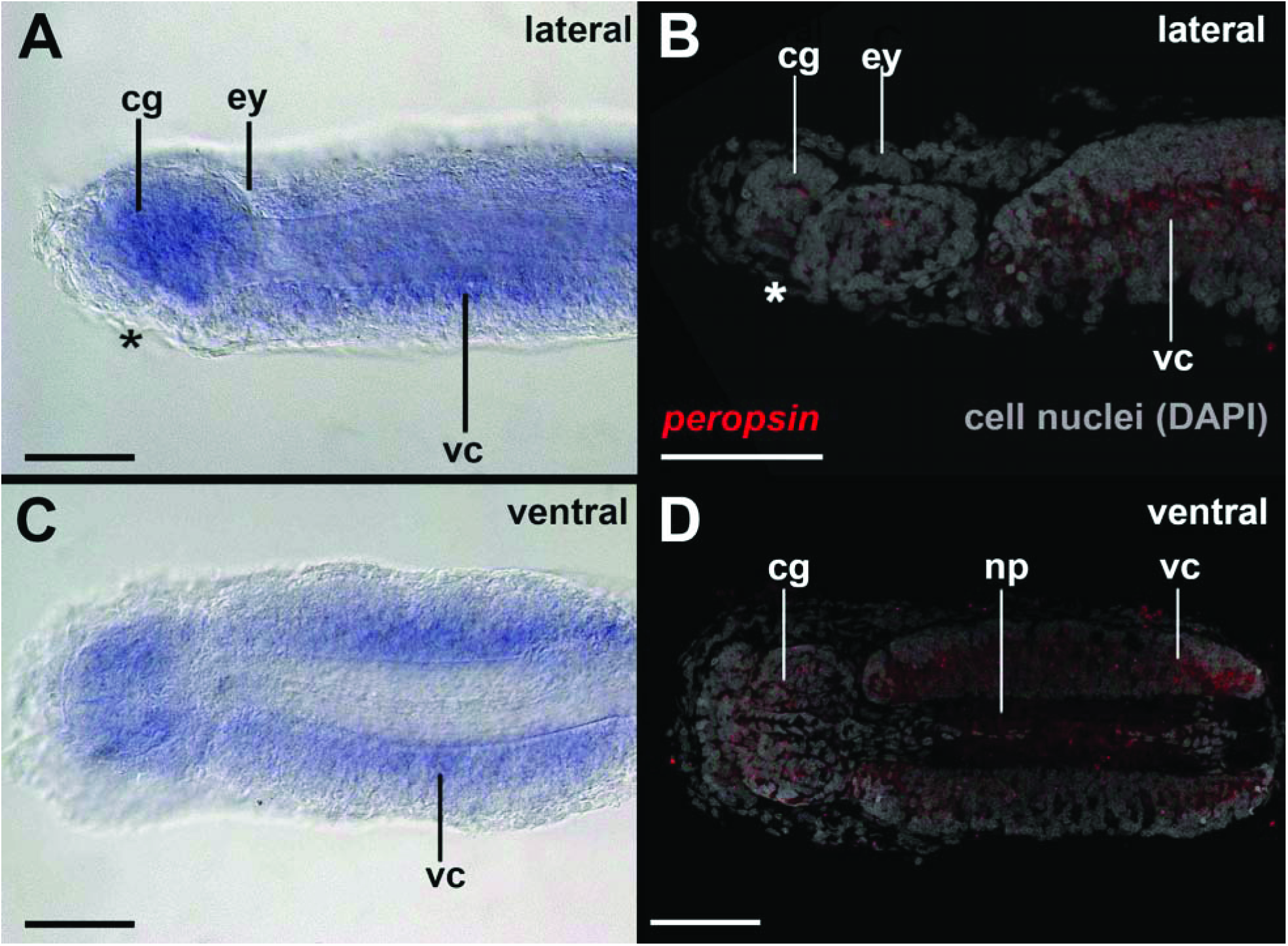
*Peropsin* expression in early hatched juveniles of *Spadella cephaloptera*. Whole-mount *in situ* hybridization of the head region, anterior to left. **A:** *Peropsin* is expressed in the region of the cerebral ganglia (cg) and the ventral nerve cord (vc) but not the eyes (ey) as highlighted by confocal laserscanning microscopy in **B. C:** Ventral view showing *peropsin*+ cells in the ventral nerve cord but not the neuropil of the latter as highlighted by confocal laserscanning microscopy in **D**. Abbreviations: asterisk, mouth; cg, cerebral ganglion; gs, grasping spines; vc, ventral nerve cord; vsg, vestibular ganglion. Scale bars: 100 µm.

**Figure 6.**
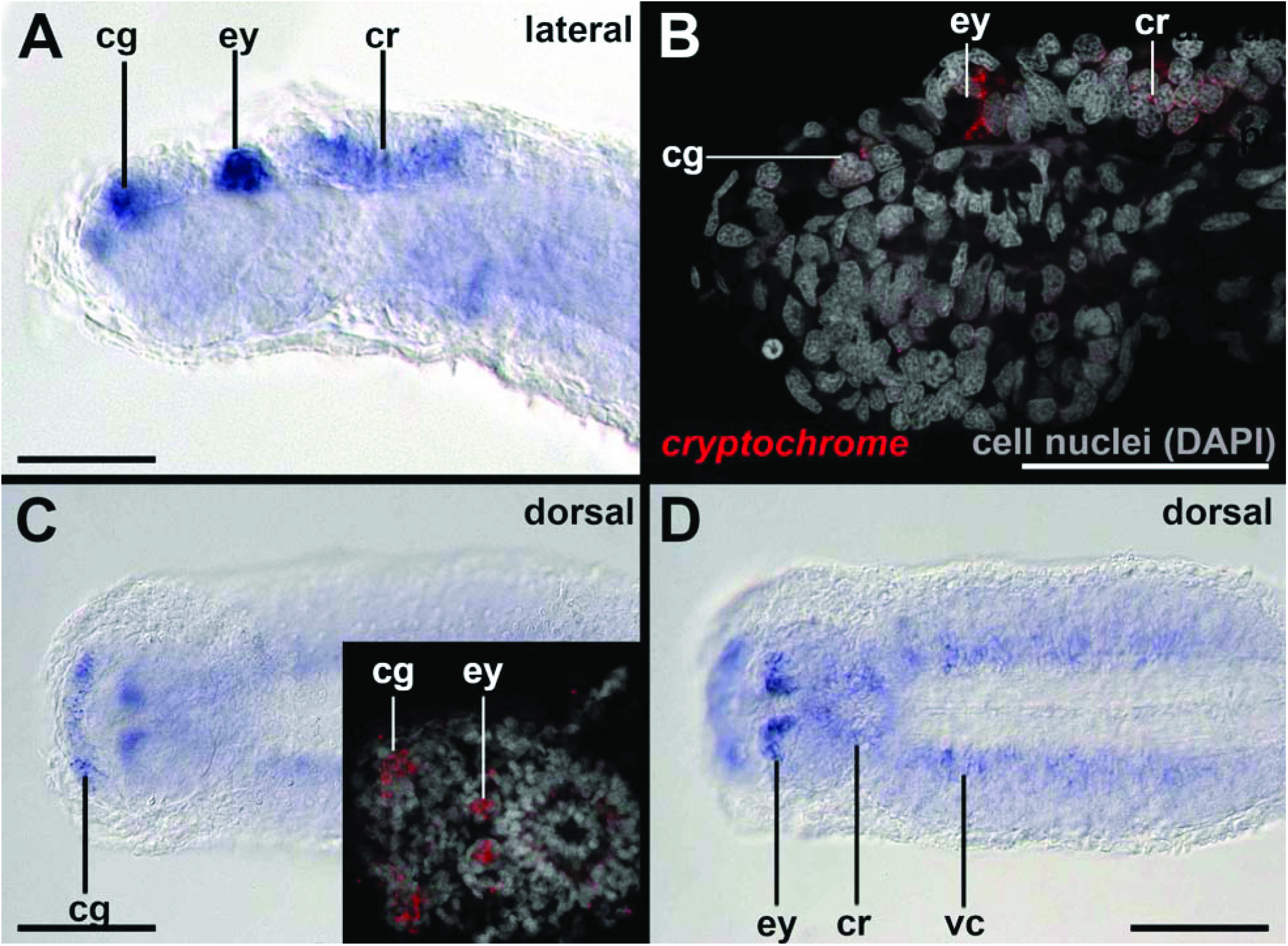
*Cryoptochrome* expression in early hatched juveniles of *Spadella cephaloptera*. Whole-mount *in situ* hybridization of the head region, anterior to left. **A:** *Cryoptochrome* is expressed in the region of the cerebral ganglia (cg), the eyes (ey), and the corona ciliata (cr) as highlighted by confocal laserscanning microscopy in **B. C:** Strong expression of *cryptochrome* in the cerebral ganglia and the proximal portions of the eyes. **D**. *Cryptochrome* expression in the ventral nerve cords (vc). Scale bars: 100 µm.

**Figure 7.**
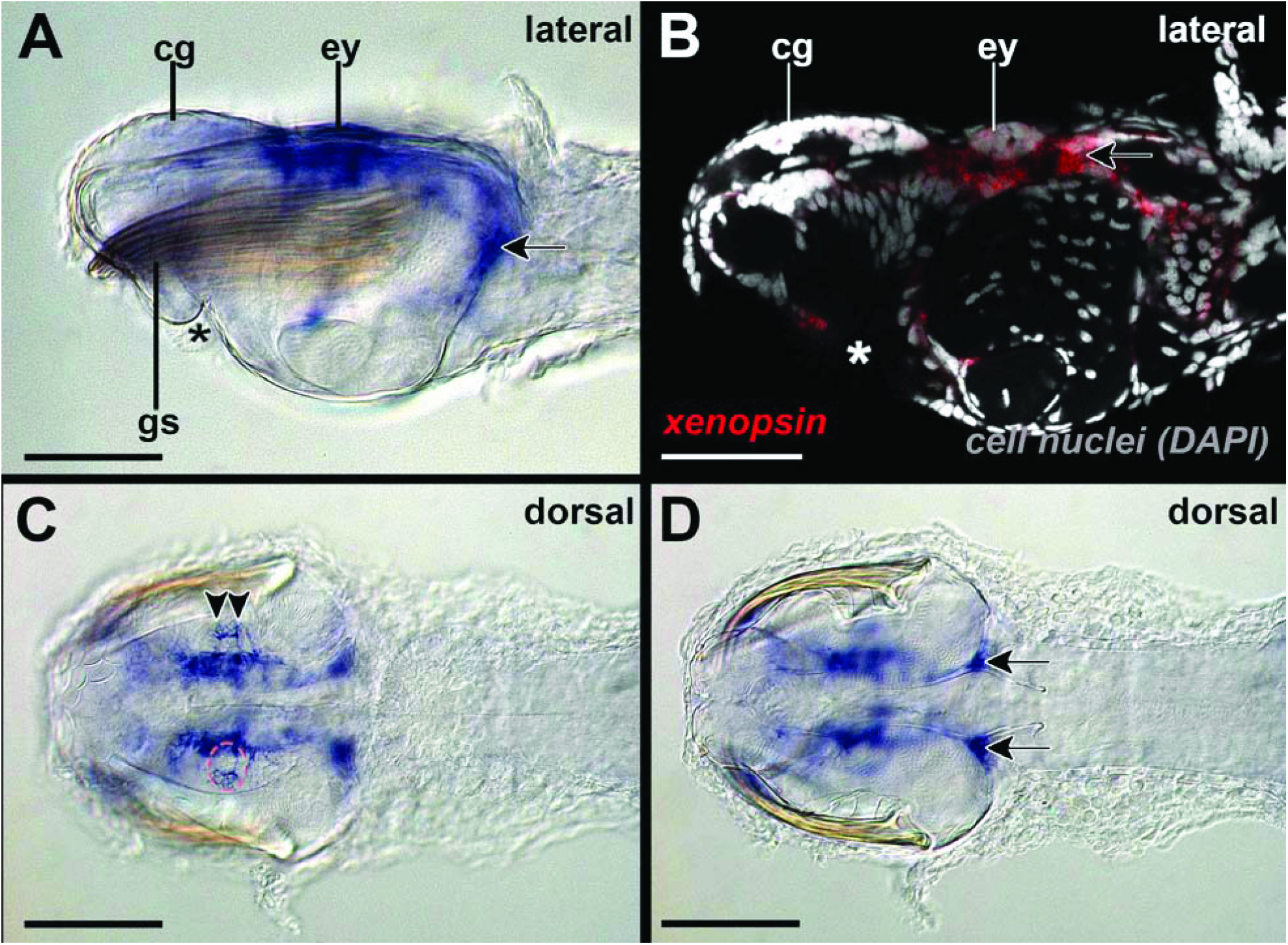
*Xenopsin* expression in adults of *Spadella cephaloptera*. Whole-mount *in situ* hybridization of the head region with anterior facing to the left. **A:** *Xenopsin* is expressed in the eyes (ey) and the region surrounding the latter. Unspecific staining marked with arrow. **B:** Confocal laserscanning reflection scan highlights *xenopsin* expression in the lateral eye and expression ventrally, anterior and posterior to it. **C:** *Xenopsin* is expressed in two photoreceptors (arrowheads) of each distolateral eye (ey) only. Stippled red circle highlights left eye. **D:** Unspecific staining marked with arrows in ventroposterior head region. Abbreviations: asterisk, mouth; cg, cerebral ganglion; gs, grasping spines. Scale bars: 200 µm.

**Figure 8.**
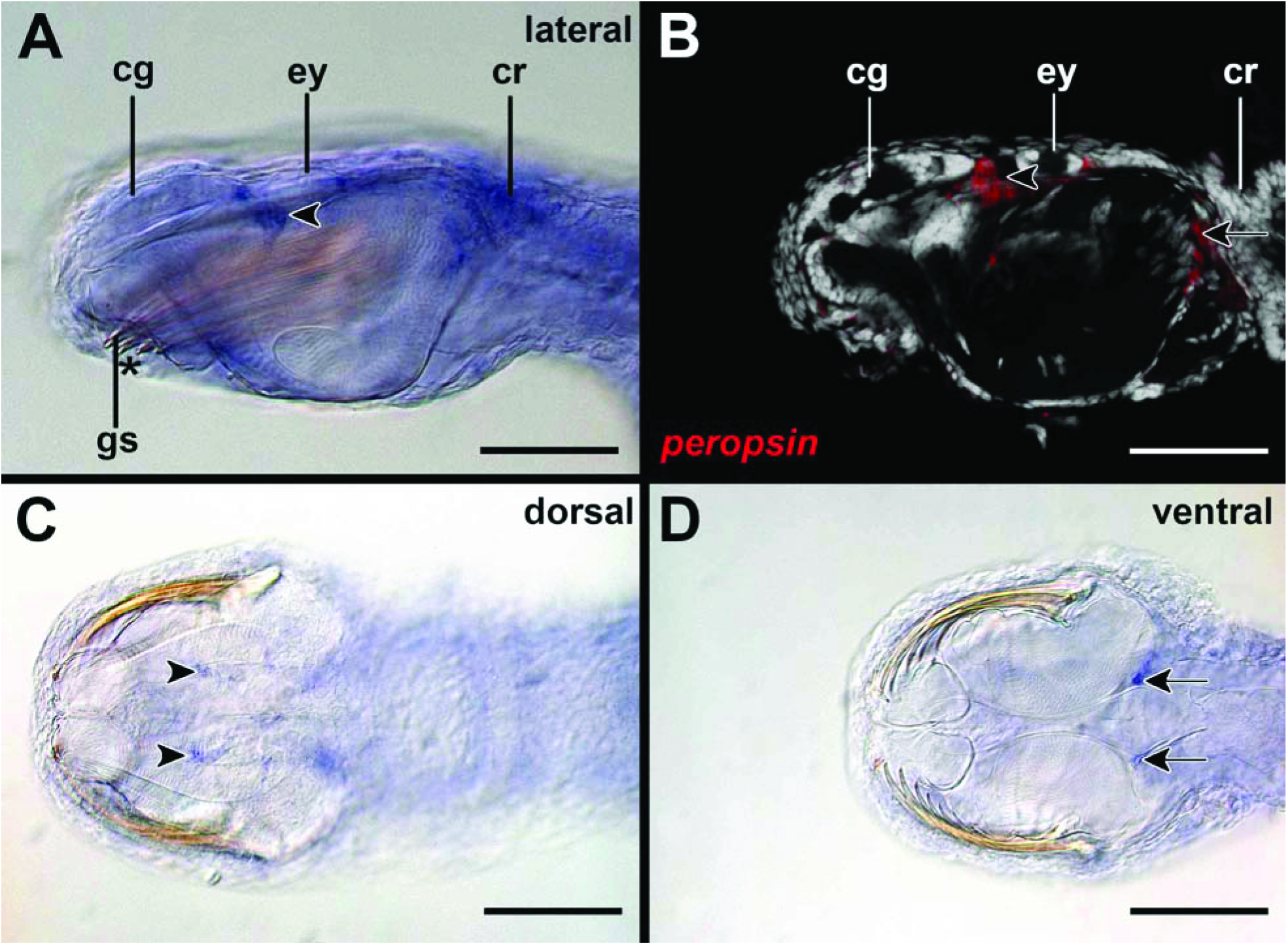
*Peropsin* expression in adults of *Spadella cephaloptera*. Whole-mount *in situ* hybridization of the head region, anterior to the left. **A:** *Peropsin* is expressed in the region (arrowhead) antero-ventrally to each eye (ey), and in the corona ciliata (cr). **B:** *Peropsin* expression antero-ventrally to the left eye highlighted by confocal laserscanning microscopy. **C:** *Peropsin* expression antero-ventrally to each eye and unspecifically stained structures (arrows; also shown in **D)**. Abbreviations: asterisk, mouth; cg, cerebral ganglion; cr, corona ciliata; gs, grasping spines. Scale bars: 200 µm.

**Figure 9.**
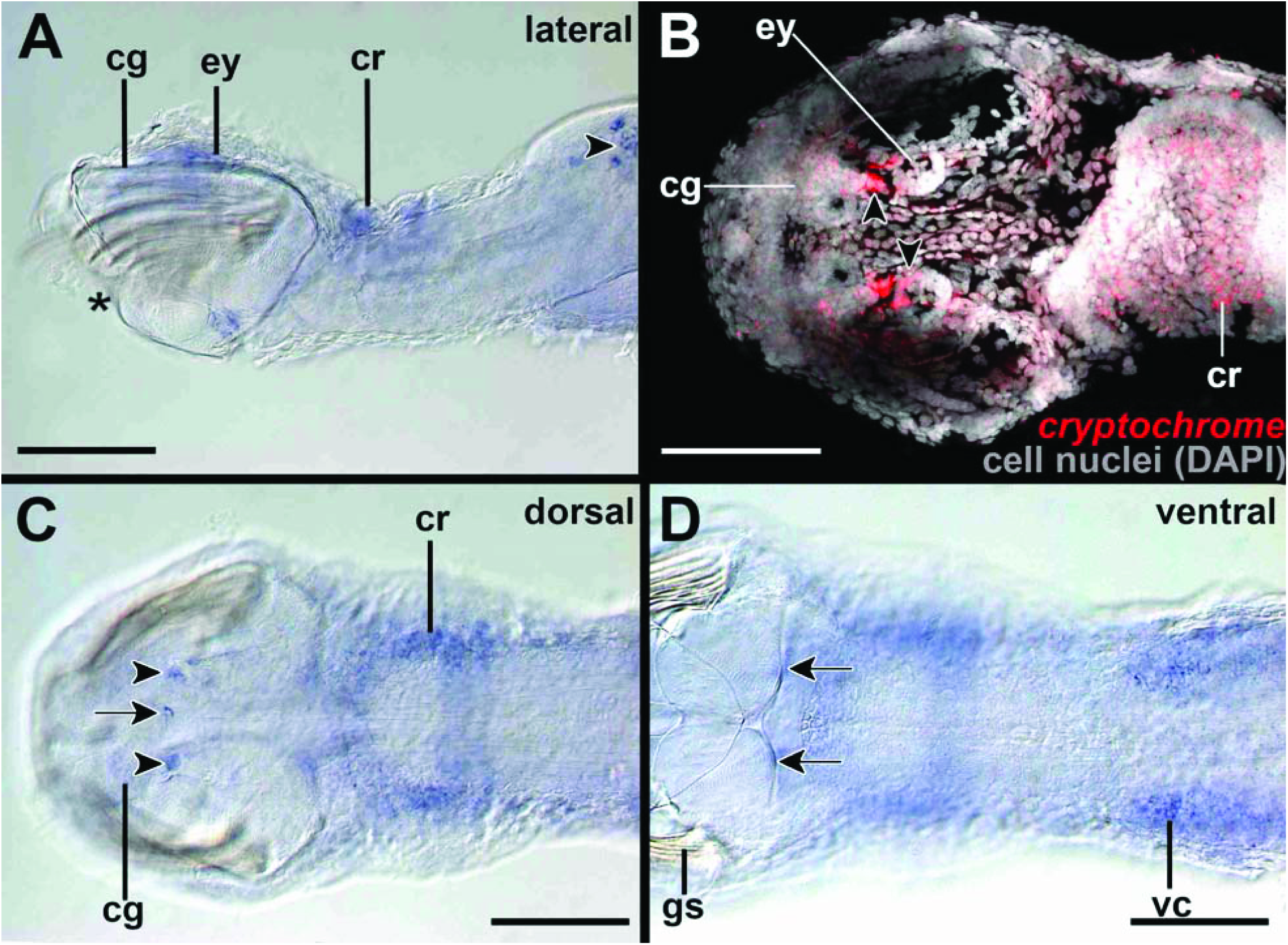
*Cryoptochrome* expression in adults of *Spadella cephaloptera*. Whole-mount *in situ* hybridization of the head region, anterior to the left. **A:** *Cryptochrome* is expressed in the region of antero-ventrally to each eye (eye) in this two weeks old juvenile. In addition, *cryptochrome* is expressed in the cerebral ganglia (cg), the corona ciliata (cr), and the few cells of the dorsal epidermis of the trunk. **B:** Adults express *cryptochrome* in the same regions but not in the dorsal epidermis as highlighted by confocal laserscanning microscopy. The *cryptochrome*+ cells are located anteroventrally to each eye (arrowheads). **C & D:** An additional *cryptochrome*+ cell (arrow) is located posterior to the retrocerebral pore, in the region between both above-mentioned expression domains (arrowheads) and in the ventral nerve cord (vc) (arrow). Unspecifically stained structures (arrows in **D**). Abbreviations: asterisk, mouth; gs, grasping spines. Scale bars: 200 µm.

## Discussion

### Ocular photoreceptors of chaetognaths express probably only a single opsin

The existence of numerous elaborate photoreceptors in the eyes suggests that vision plays an important role during predation and escape responses of chaetognaths (Eakin and Westfall 1964; Goto and Yoshida 1984). Our thorough screen of a transcriptome derived from total RNA of various developmental stages and adults of *Spadella cephaloptera* however only yielded single-copy orthologs of *sce-xenopsin* and of *sce-peropsin*. Note that two copies of *xenopsin* have been identified for another chaetognath species, *Pterosagitta draco* (Rawlinson et al. 2019). Our subsequent gene expression analysis demonstrated that *sce-xenopsin* but not *sce-peropsin* is expressed in the eyes of developmental stages and adults of the chaetognath *S. cephaloptera* (Figs. 3-5). In early embryos and hatchlings *xenopsin*+ photoreceptors are located in the lateral portions of the eyes (Fig. 4C-D), raising the question of whether the anterior and posterior cells of the eyes are indeed photoreceptors and if so, which opsin is expressed in the latter. To answer this question, an intensive search for other light-sensitive proteins identified an ortholog of *cryptochrome. Sce-cryptochrome* is expressed in fewer cells than *sce-xenopsin* but in the same lateral domains in hatchlings (although not in adults) (Figs. 4, 6, 7, 9). Co-expression of *sce-xenopsin* and *sce-cryptochrome* in the lateral photoreceptors of the chaetognath eyes is reminiscent of the condition in the annelid *Platynereis dumerilii* in which *cryptochrome* and *c-opsin* are co-expressed in the apical nervous system (Tosches et al. 2014).

### Non-visual light sensitive cells in the chaetognath nervous system?

*Sce-cryptochrome* and *sce-peropsin* are both expressed in the dorsal cerebral ganglia and the ventral nerve cord, a condition that resembles the co-expression of *cryptochrome* and *peropsin* in the apical nervous system of the annelid *Platynereis dumerilii* (Tosches et al. 2014). These findings may indicate that the dorsal cerebral ganglia and portions of the ventral nerve cord are domains of non-visual light sensitivity as has been proposed for *P. dumerilii*, in which light detection, circadian entrainment and melatonin release are performed by a single photoreceptive zone (Tosches et al. 2014).

### The corona ciliata, an organ with light-sensitive receptors?

Many different roles have been proposed for the corona ciliata including excretory (chaetognaths lack nephridia), (chemo)sensory, or secretory (glandular) function (Shinn 1997; Müller et al. 2014; Harzsch et al. 2015). Our study demonstrates that *sce-cryptochrome* is expressed in distal and proximal cells of the corona ciliata suggesting that cells of this organ may be involved in circadian entrainment (Fig. 9A-C).

### The Spiralia – a clade unveiling tremendous photoreceptor diversity

Recent studies revealed diverse combinations of photoreceptors and opsins within the Spiralia (Passamaneck et al., 2011; Vöcking et al. 2017; Rawlinson et al. 2019). Trochophore larvae of the polyplacophoran mollusk *Leptochiton asellus* exhibit multi-ciliated and microvillar photoreceptors that co-express *xenopsin* and *r-opsin*, while the larva of the brachiopod *Terebratalia transversa* exhibits eyespots with ciliated photoreceptors expressing *xenopsin* (Passamaneck et al. 2011). A recent study on the platyhelminth *Maritigrella crozieri* showed *xenopsin* expression in the larval epidermal eye, in the cerebral eyes, and in the adult phaosomal photoreceptors, which all consist of ciliated photoreceptors (Rawlinson et al., 2019). In the larval epidermal eye of *M. crozieri*, a single photoreceptor cell produces many cilia which form a lamella packed into a pigmented pocket formed of an adjacent cell, similar to how the array of chaetognath ciliated photoreceptors jut into one of several pockets formed in a central pigment cell (Rawlinson et al. 2019; Fig. 2C, D). Cilia of the chaetognath photoreceptors elaborate into annulated lamellae within these pockets (Goto et al. 1984). While the chaetognath eye is not phaosomous as described earlier (Purschke et al. 2006), it is encapsulated by the extracellular matrix and sheath cells as are the phaosome and epidermal eye of the platyhelminth *M. crozieri* (Rawlinson et al. 2019). It has been speculated that the phaosomal photoreceptive structures of platyhelminths are evolutionarily derived (Sopott-Ehlers et al., 2001), however, they are also present among annelids and phaosome-bearing annelids have not been exhaustively searched for *xenopsin* orthologs yet. Hence, it is tempting to speculate that each species-specific phaosome-like structure may be derived from a ciliated photoreceptor expressing *xenopsin* that was present in the last common ancestor of Spiralia. Together, our and other comparative studies emphasize the importance to investigate taxa such as Gnathifera to unravel the evolution of spiralian and bilaterian body plans.

## Methods

### Ethics, collection and culture of animals

Individuals of the chaetognath *Spadella cephaloptera* (Busch, 1851) were collected in front of the Station Biologique de Roscoff, Roscoff, France in summer 2018 and transferred to an aquarium at the European Molecular Biology Laboratory (EMBL) in Heidelberg. Chaetognath are hermaphrodites and adults started to reproduce in the aquarium at 18°C water temperature. The date of egg deposition and hatching was documented with an accuracy of 12 hours.

### RNA extraction and fixation of animals for *in situ* hybridization experiments

Adult *Spadella cephaloptera* were starved for three days and sacrificed together with developmental stages covering early zygotes to hatched 1 months old juveniles for RNA extraction by using a RNA extraction kit (Qiagen, Roermond, Netherlands). Additional adults and developmental stages were carefully anesthetized in 7.14% MgCl_2_ before fixation and fixed for 1 hour at room temperature for *in situ* hybridization experiments and treated as previously described (Wollesen et al. 2015). Extracted RNA was used for transcriptome sequencing (see below) and for the synthesis of cDNA used for subsequent riboprobe synthesis.

### Transcriptome sequencing and assembly

For the transcriptome pooled total RNA was Illumina 150bp paired end sequenced and resulting in a total of 57,486,532 paired reads. The short-read libraries were pre-processed using trimmomatic (v. 0.36; Bolger et al. 2014) to remove known specific Illumina adapters from the paired-end libraries (Illumina universal adapter). Filtering by quality and length was performed with a SLIDINGWINDOW:4:15 MINLEN:36. First and last nucleotides from reads with low quality score were clipped and the library file was converted into fasta format using fq2fa from SeqKit (version 0.11.0). Quality of the initial and filtered library was assessed with the software FastQC (v.0.11.8; Wingett and Andrews 2018) considering quality score of the bases, GC-content, and read length. 11.83% of reads were excluded during the pre-processing procedure resulting in a total of 50,686,453 reads. The assemblies and all downstream analyses were conducted with a high-quality and clean library. The filtered transcriptome was assembled into contiguous cDNA sequences with IDBA_tran v1.1.3 software (Peng et al. 2013) using the default settings (except: -mink 20 -maxk 80 -step5). The resulting assembly was assessed using the tool QUAST (http://quast.bioinf.spbau.ru). The number of contigs was 148988 with 8.4137 contigs longer than 1000 bp. The number of reconstructed bases was 259.028.025 with 222.358.174 contigs longer than 1000 bp. The length of the largest contig was 35.950,00, the N50 2.179,17, the N75 1.592,00 and the GC content 45,74%.

### Alignment and phylogenetic analysis

Candidate genes were identified by BLAST searches of bilaterian orthologs against the transcriptome of *Spadella cephaloptera* (see above). The phylogenetic analysis was performed for the predicted protein sequences of Sce-Xenopsin and Sce-Peropsin building upon the analyses of Vöcking et al. (2015, 2017) and Ramirez et al. (2016). Orthologous sequences from various metazoan species were downloaded from NCBI (https://www.ncbi.nlm.nih.gov/) and Uniprot (https://www.uniprot.org). Multiple sequence alignment was performed with MAFFT v7.123b (Katoh & Standley 2013) and the alignment was manually trimmed in AliView v.1.26 (Larsson A 2014). A bayesian phylogenetic analysis on 71 taxa and 364 characters was carried out with MrBayes v3.2.6 in CIPRES Science Gateway (https://www.phylo.org). The LG+G amino acid replacement model was estimated with Prottest3 v3.4.2 (Darriba et al. 2011), in addition to 3245000 generations. The resulting consensus tree was visualized and adjusted in iTOL v4.4.1 (https://itol.embl.de/about.cgi). For the phylogenetic analysis of cryptochromes the overall procedure was the same with exceptions. Sequences used in the phylogenetic tree by Ozturk (2017) were included in this phylogenetic tree. A bayesian phylogenetic analysis on 37 taxa and 510 characters was carried out with MrBayes on XSEDE v3.2.7a in CIPRES Science Gateway (https://www.phylo.org). The LG G+I amino acid replacement model was estimated with Prottest3 v3.4.2 (Darriba et al. 2011), in addition to 205000 generations until the average standard deviation of split frequencies got <0.01 (0.009943). The resulting consensus tree was visualized and adjusted in iTOL v4.4.1 (https://itol.embl.de/about.cgi).

### Molecular isolation of RNA transcripts

A first strand cDNA Synthesis Kit for rt-PCR (Roche Diagnostics GmbH, Mannheim, Germany) was used for first-strand cDNA synthesis of the RNA pooled from different developmental stages of *Spadella cephaloptera.* Identified gene sequences in sense orientation were used to design gene-specific primers with an annealing temperature of >60°C. Reverse primers containing part of the T7 promotor sequence (5’-TAA TAC GAC TCA CTA TAG GG-3’) followed by a reverse complement of the gene specific sequence. PCRs were successfully carried out with the following primer sequences:

*sce_xenopsin*

Ffw GTCGGACTTGTTCATGTGCTCCGTC

Rev TAATACGACTCACTATAGGGCGTCGCGGTGACGCTCAGTTC

*sce_peropsin*

Ffw CTGGATCTGGGGAGATCTCGGCTG

Rev TAATACGACTCACTATAGGGGAGCCTCGGGAAATCGGAGACCAC

*sce_cryptochrome*

Ffw GATCCTCGGGAGATCTTTGCTCGTCTC

Rev TAATACGACTCACTATAGGGGCCAAAGATGACCGCGCGTGAG

5µL of each PCR product were size-fractioned by gel electrophoresis and the remaining 45µL of the PCR product were cleaned up with a QIAquick PCR Purification Kit (QIAgen, Hilden, Germany) if size estimation and expected sequence length matched. PCR products were sent for sequencing to confirm gene identity and sequences of *sce-xenopsin, sce-peropsin*, and *sce-cryptochrome* were deposited on Genbank (Accession numbers: Sce_xenopsin (MN735184), Sce_peropsin (MN735185), Sce_cryptochrome (MN735186)).

### Probe synthesis and whole-mount *in situ* hybridization

*In vitro* transcription reactions were performed with the above-mentioned templates, digoxigenin-UTP (DIG RNA Labeling Kit, Roche Diagnostics), and T7 polymerase (Roche Diagnostics GmbH) for the synthesis of antisense riboprobes, according to the manufacturer’s instructions. For whole-mount *in situ* hybridization experiments, specimens were rehydrated into PBT (phosphate buffered saline + 0.1% Tween-20) and treated with Proteinase-K at 37°C for 10 min (10 µg/ml in PBT). Specimens were pre-hybridized in hybridization buffer for 4-10 h at 63°C (see Wollesen et al. 2015 for details). Hybridization was performed at the same temperature with probe concentrations ranging between 1-2 μg/ ml for 21-24 h. A DIG-labeled AP-antibody was used at a dilution of 1:2500 in blocking solution at 4°C overnight. Color development in the NBT/ BCIP/ Alkaline Phosphatase buffer solution took 6-24 hrs at 4°C. Some specimens were counterstained with DAPI to visualize cell nuclei (Sigma-Aldrich; St. Louis, MO, USA). A minimum of 30 individuals per stage were investigated. The majority of whole-mount preparations were cleared in a solution of 2,2’thiodiethanol (Sigma-Aldrich), mounted on objective slides and analyzed. Preparations were documented with an Olympus BX53 Microscope (Olympus, Hamburg, Germany). In addition, developmental stages were scanned with a Leica confocal SP8 microscope (Leica Microsystems, Wetzlar, Germany) using brightfield, autofluorescence, and reflection mode scans to document the precise cellular location of transcripts (Jékely and Arendt 2007). If necessary, images were processed with GIMP (version 2.10.10, www.gimp.org) to adjust for contrast and brightness. Sketch drawings were created with Inkscape (version 0.92.4, www.inkscape.org).

## Abbreviations

*Sce*: *Spadella cephaloptera*;
BCIP: 5-brom-4-chlor-3-indoxylphosphat;
BLAST: Basic local alignment search tool;
cDNA: complementary deoxyribonucleic acid;
DIG: digoxigenin;
NBT: nitro blue tetrazolium;
NCBI: National center for biotechnology information;
PBT: Phosphate buffered saline with tritonX-100;
pcr: polymerase chain reaction.

## Authors’ contributions

TW designed the study, raised the chaetognaths, extracted the RNA, and carried out all experiments. SVR and TW assembled the transcriptome and carried out the phylogenetic analyses. TW drafted the manuscript and SVR, APO, and DA commented on the manuscript. All authors read and approved a final version of the manuscript.

## Acknowledgements

TW thanks Jacob Musser (EMBL) for submitting RNA for transcriptome sequencing. The EMBL GeneCore facility is thank for transcriptome sequencing.

## Availability of data and materials

All sequences analyzed in this study have been published on publicly accessible websites.

## Ethics approval and consent to participate

Not applicable.

## Consent for publication

Not applicable.

## Figure legends

**Supplementary Figure 1.**
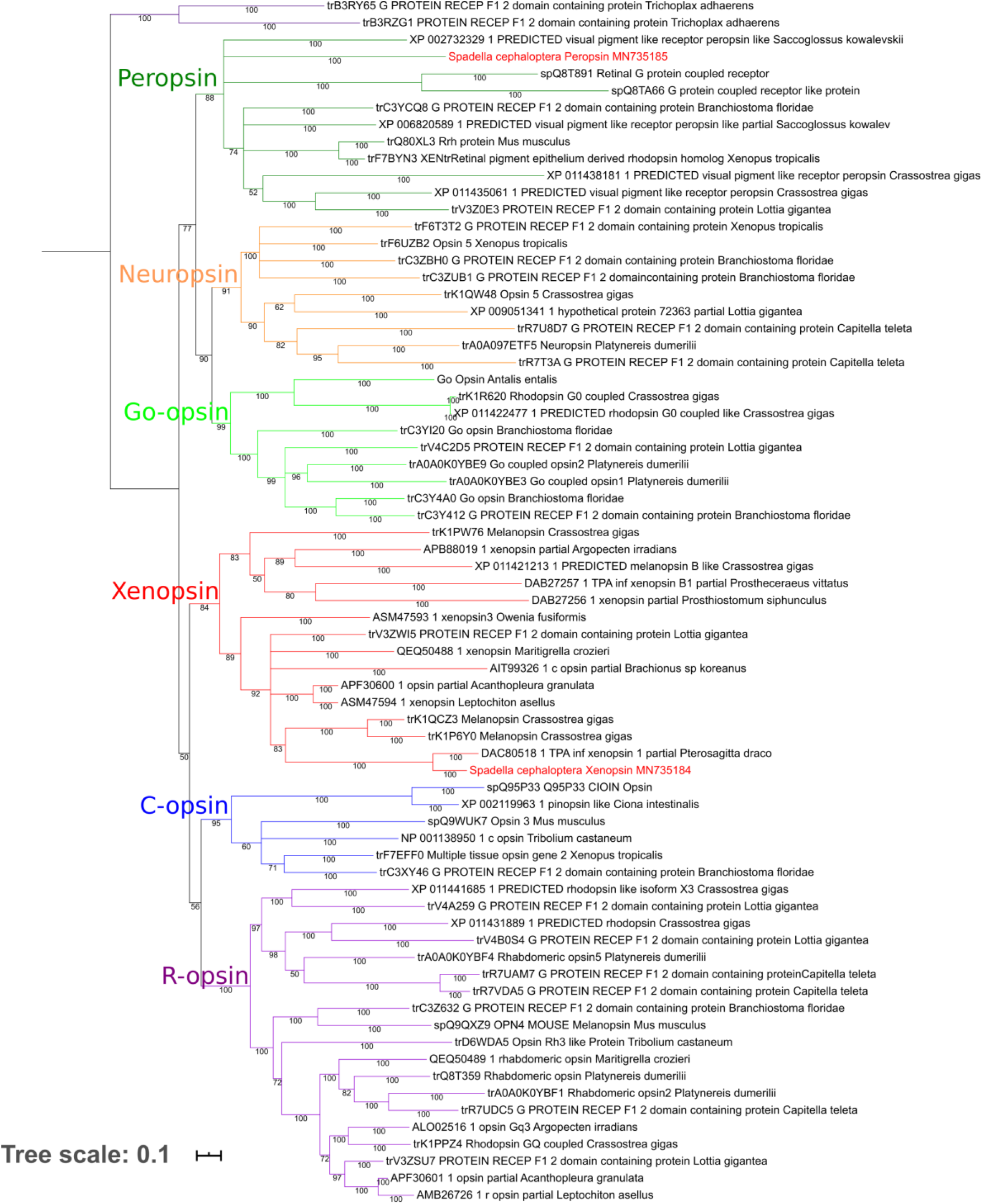
Phylogenetic tree of opsins. Majority consensus phylogenetic tree based on Bayesian analysis. Bootstrap values are shown. Xenopsin and Peropsin sequences of the chaetognath *Spadella cephaloptera* are highlighted in red. Nomenclature follows Ramirez et al. 2016. The Placopsins of *Trichoplax adhaerens* served as outgroups.

**Supplementary Figure 2.**
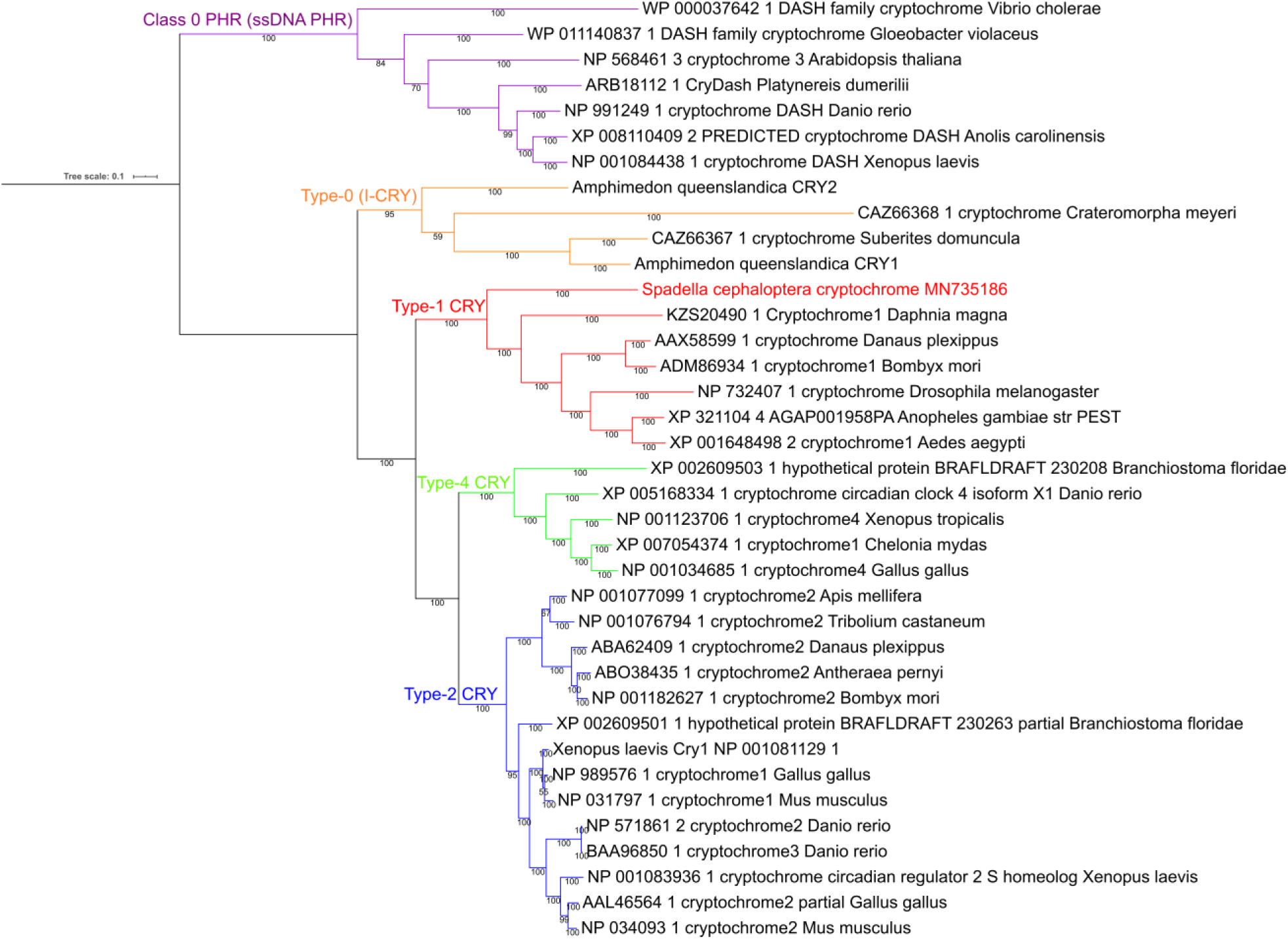
Phylogenetic tree of cryptochromes. Majority consensus phylogenetic tree based on Bayesian analysis. Bootstrap values are shown. The Cryptochrome of the chaetognath *Spadella cephaloptera* is highlighted in red. Nomenclature follows Ozturk (2017). Class 0 PHR (ssDNA PHR) cryptochromes serve as an outgroup.

